# Omics and bioinformatics approaches to target boar taint

**DOI:** 10.1101/079301

**Authors:** Rahul Agarwal, Jitendra Narayan

## Abstract

In livestock species, a rapid growth in high-throughput omics data has accelerated the pace of studies that target to dissect economically important traits to provide better quality animal products to consumers. In pig industries, young boars are generally castrated to remove boar taint, a phenotypic and inheritable trait well-known by an abnormally bad smell and taste in pork meat derived from some uncastrated male pigs. Existence of porcine reference genome made possible to catalogue genome-wide QTLs, candidate genes and biomarkers in associations with boar taint and other industrially significant traits in pigs. The aim of this paper to review the contribution of bioinformatics resources and omics technology in boar taint related studies. This paper also provides concise details about state-of-the-art sequencing technology.

## Introduction

There are economic limitations in raising castrated boars for pig breeders since they entail supplementary feed and have not a satisfactory meat to fat ratio. Boars usually castrated when they reach puberty to avoid the incidence of boar taint, an unpleasant smell and taste in uncastrated boars meat (Bonneau M, 1997). Contrary, castration also results in loss of sex steroids from male pigs, which in turn affects the growth of pigs and economically, obtained meats are not of desirable quality as what consumers expect. These serious implications push EU and some EEA countries to implement ban castration of male pigs by 2018. Surgical castration is routinely performed by pig industries to eradicate androstenone and skatole (Patterson 1968; Vold 1970), the two major chemical compounds whose excess deposition in adipose tissue accountable for boar taint in some mature male pigs. Androstenone (5α-androst-16-en-3-one) is a 16-steroid compound synthesized in the Leydig cells of the testis (Patterson R, 1968) while skatole (3-methylindole) yielded, as a result of microbial breakdown of tryptophan in colon part of male and female pigs. Androstenone blocks skatole metabolism (Doran, Whittington et al. 2002; Chen, Cue et al. 2008;Rasmussen 2011), which pointing out that an elevated levels of skatole is merely an issue in male pigs. Besides, indole, 2-aminoacetophenone (a hepatic skatole metabolite) and androstenol are other compounds too contribute toward incidence of boar taint though with a lesser extent (Hansson, Lundstrom et al. 1980; Jensen, Cox et al. 1995). High genetic correlations to testosterone and estrogens (0.8-0.9 for androstenone and 0.4-0.6 for skatole for the Norwegian breeds) are though a major obstacle as there is no clear answer to how selection would affect fertility of the pigs. Hence there is need to formulate alternate solution to remove boar taint without comprising the reproduction levels. Since storage of both androstenone and skatole in adipose tissue is highly heritable, therefore, it is conceivable genetically to select the boars with reduced concentration of androstenone and skatole. Genetic selection can be the most promising alternative solution as suggested by (Fredriksen and Nafstad 2003), unfortunately owing to the large number of genes affecting boar taint as well as strong correlations to steroid hormones confer hindrance to its implementation. However, this approach requires the integration of bioinformatics, all-omics, and genetics.

Owing to the mountainous growth in the amount of biological data generated from different omics technologies in recent times, the involvement of bioinformatics becomes indispensable. Now, it is feasible to detect innumerable number of putative variants (SNPs, small InDeLs, copy number variations and structural variations) from the billions of raw short-reads generated by sequencing the multiple genomes, exomes or transcriptomes of an organism simultaneously using latest 2^nd^ and 3^rd^ generation sequencing machines. Particularly, SNPs (single nucleotide polymorphisms) that are present abundantly in eukaryotic genomes could be useful for tracing out the entire candidate genes, mutations and alleles found associated with boar taint using association study. Consequently, these SNPs, which either individually or in combinations (haplotypes) significantly linked with boar taint compounds, can be utilized for genetic selection of boars with a minimal amount of boar taint levels.

## Introduction to Sequencing

An era of sequencing started with the arrival of two sequencing methods in 1970s namely Maxam-Gilbert sequencing (Maxam et al., 1977) and Sanger sequencing (Sanger et al., 1977). Maxim-Gilbert sequencing, a commonly known as chemical based sequencing in which the DNA fragment chemically modified at 5’ end and consequent partial cleavage of fragment takes place at sites adjoining to modified bases. Later on, the Sanger sequencing method, alternatively known as a chain - termination method based on the use of both terminator dideoxynucleotides (ddNTP’s) and standard nucleotides (NTP’s) naturally found in DNA, developed by Frederick Sanger has been popular for almost two decades after earlier method failed to capitalize due to limitations in terms of time and use of toxic chemicals. Completion of the Human Genome Project (HGP) (Lander et al., 2001; Venter et al., 2001) by Sanger sequencing is one of the major attainments in sequencing history. This project took almost 13 years to complete and costs billions of US dollars. The major disadvantage of the Sanger method is its lack of ability to parallelize though longer reads generated by this method (Shendure, 2008; França et al., 2002). However, there has been quantum shift happened in sequencing technology in past decade, now scientists can imagine to sequence entire human genome under 1000 US dollars with very recent launch of Illumina Hi-Seq X Ten, an integrated system of ten Hi-Seq X sequencers capable to sequence above 18,000 genomes per year. The per-base price of DNA sequencing has plunged drastically since year when HGP accomplished and now surpassing the expectation of Moore’s law by many miles.

Many commercial sequencing companies such as Illumina, Life Technologies, Pacific biosciences, and Oxford Nanopore (Lin et al., 2012; Pareek et al, 2011) are in the race to deliver a efficient and cost-effective sequencing machine to the scientific fraternity, biotechnology and pharmaceutical companies. Current sequencing machines (França et al., 2002) are classified on the basis of following methods- sequencing by hybridization, pyrosequencing, sequencing by synthesis, sequencing by ligation and single molecule sequencing. HiSeq 2000 (Illumina Inc.,2010)is capable of generating 600Gb data in approximately 11 days and read length of 100bp is now one of the most widely used sequencers and its chemistry of obtaining reads derived from Solexa sequencing. Illumina sequencing method (Kircher et al., 2011; Ansorge, 2009) ensues following two processes sequentially:

### 1) Clonal amplification

First, randomly fragmented PCR enriched DNA molecules ligated with adaptors at both ends are attach to the solid surface of the flow cell. The flow cells attached the single stranded DNA fragments in such a way that the access of enzymes will not be affected and at the same time ensure the stability of the fragments bound to the surface. The flow cells already enriched with primers. Then 3’end of DNA fragments that bound to the complementary primers initiate the process of solid-phase bridge amplification. In this particular type of amplification, unlabeled dNTPs and enzyme used to extend the single stranded DNA fragment to form a double stranded molecule in which an immobilized part of double stranded molecules flip over to bind with complementary adjacent primers to form a bridge while its mobilized part washed away. This process repeated multiple times to generate hundreds of millions of dense clusters of input fragments (Shendure, 2008).

### 2) Sequencing by synthesis and cyclic reversible termination

The forward strand of each cluster is hybridized with primers to initiate the sequencing cycle in which each primed strand gets extended by DNA polymerase through incorporation of radioactively labeled dNTPs. Incorporation of the particular base next to the primer and so on is decided on the basis of signal intensity by the laser. Sequencing is a cyclic process so for each position; the signal intensities of four bases estimated multiple times that minimize the raw base calling error. Each incorporated dNTPs acts as terminators (Metzker et al., 1994) which block the extension of the cluster until imaging and cleavage step finished. In cleavage step, there is the removal of a terminating group of dNTP and fluorescent dye to continue the extension process (Metzker, 2010).

The signal intensity per cycle and signal intensity per position are plotted to decode the raw base values into a sequence of A, C, G and T (Ledergerber et al, 2011; Suzuki et al., 2011). Each read sequence is usually put together with its quality score in text based format called as *fastq file* with file extension either *.fq* or *.fastq* (Cock et al., 2010). The base quality score (Ewing et al., 1998; Ewing and Green, 1998) is used to estimate the accuracy of each base in read sequence. Each base is assigned a phred like quality score (Q), which is the probability that the sequenced base wrongly called. The higher the quality score value of the base, the higher the chance of that base being called accurately. For example, a quality score of 30 (Q30) indicates an error rate of 1 in 1000, with a related call accuracy of 99.9%. The base quality score is found to be a show decreasing trend toward the 3’end of the reads and also depend on the length of the reads (Smith et al., 2008). Low base quality score can create a problem in calling variants (Brockman et al., 2008). Therefore, it is important to eradicate those bases that have a low Q score.

## Pig Genome and its application

The proposal of pig genome sequencing started with the foundation of Swine Genome Sequencing Consortium (SGSC) in 2003 (Archibald et al., 2010; Schook et al., 2005) with an aim to provide the pig reference genome to scientific communities around the world to accelerate biomedical and health related studies including investigation of multifactorial diseases such as obesity, diabetes, dyslexia and Parkinson’s disease (Lunney, Joan K, 2007,Walters et al., 2012). A physical map of the pig genome was created with utilization of 267,884 numbers of restriction fragments, and over 600k BAC end sequences (BES) from four large insert bacterial artificial chromosome (BAC) clone libraries. These BES sequences are available at http://www.ncbi.nlm.nih.gov/Traces/trace.cgi. Chromosome-wise pig clone libraries can be accessed at https://www.sanger.ac.uk/cgi-bin/humace/clone_status. Result of alignment between BES sequences and human genome using Wu-Blastn tool was integrated with data of fingerprints and restriction hybrid (RH) markers to construct map with 172 contigs using WebFPC tool (http://www.agcol.arizona.edu/software/fpc/) and distributed across 18 autosomes and 2 sex chromosomes X and Y. Subsequently, the map (http://www.marc.usda.gov/genome/swine/swine.html) was used to select minimal tiling path (MTP) across the pig genome for clone based sequencing to provide initial draft with 4-6x genome coverage using phrap (http://www.phrap.org/) and SSAHA tools. BAC clones based sequences combined with Illumina/Solexa based data generated from whole genome shotgun sequencing of a DNA collected from a single female Duroc breed (Duroc 2-14) using BLAT alignment tool (Humphray et al., 2007; Uenishi et al., 2012; Groenen et al., 2012). Assembly of whole genome data was done with help of SOAPdenovo and Cortex genome assemblers.

There were many drafts of the pig genome released so far, and latest one is build 10.2.75 which is consists of ~3Gb bps stacked into 20 chromosomes and 4562 unplaced scaffolds. Current pig genome assembly, i.e., Sscrofa10.2 assembly is annotated with 21,640 protein coding genes, 2,965 ncRNAs and 380 pseudogenes. This assembly is stored in Genbank with its accession *GCA_000003025.4*. More information about latest build is available at http://www.ensembl.org/Sus_scrofa/Info/Index. However, this current pig reference genome may oversight a precise copy number of alleles in the original Duroc sow’s genome (Warr A et al., 2016). In addition, the Sino-Danish pig genome sequencing project was launched separately in which altogether 3.8 million shotgun sequencing reads and approximately 1 million EST sequences was generated and available to download at http://www.piggenome.dk/. Presence of whole genome data in pig would be useful for discovering more QTLs, candidate genes and haplotype data associated with economically significant traits using an abundance of high quality genetic markers.

Pig genome database (PiGenome) is a source to browse multiple details at the transcripts and genome levels and can access at http://nabc.go.kr/sgd/index.php. This database is a repository of pig ESTs (Expressed Sequence Tags), consensus sequences, BAC clones, BES (BAC End Sequences) and quantitative trait locus (QTLs). ArkDB (http://bioinformatics.roslin.ed.ac.uk/arkdb/) is another vital resource to obtain information of references, markers and loci and genetic linkage and cytogenetic maps in farmed animals including pig. Roslin Bioinformatics (http://bioinformatics.roslin.ed.ac.uk/) is a web portal to access important genetic data and various tools to perform data analysis work in pig and other species. Other sources to obtain pig related data are http://www.animalgenome.org/pig/ and URL given in table 1 of this paper (http://www.karger.com/Article/FullText/324043).

Availability of the reference genome of many eukaryotic species pushes the work of whole genome re-sequencing (Bentley, 2006; Pareek, 2011) to the tremendous pace. Whole genome re-sequencing is a technique to sequence multiple genomic DNA all together so as to provide enough coverage for calling true variants. It is now possible to perform re-sequencing of the whole genomes, whole exomes, multiple targeted regions or selected genes in several individuals concurrently. From re-sequencing data (Stratton, 2008), identification of polymorphisms, small InDeLs, copy number variation and structural variations amongst aligned reads and between sequenced reads and the reference genome can be attainable (Amaral et al., 2009; Aslam et al., 2012; Stothard et al., 2012). Nowadays, re-sequencing contributes significantly in association studies to fine map the QTLs, ascertain causative mutation and in the genetic selection of animals against a particular phenotypic trait like boar taint (Day-Williams et al., 2011). With remarkable drop in the price of sequencing, whole genome re-sequencing of thousands of individuals is increasingly popular in livestock species and invertebrates, in addition to, the 1000 Human Genomes Project, which launched some years ago to compile the list of all the rare and common genetic variations in human populations living across different continents of the world.

## Utilization of omics technology in boar taint

### DNA sequencing

In 2009, high density and high- throughput assay of 64,232 SNPs was constructed on Illumina platform and known by commercial name ‘PorcineSNP60’. These SNPs detected from whole genome sequencing data of four pig breeds (Duroc, Landrace, Large White, Pietrain) and a wild boar population and sequencing performed on 1G Genome Analyzer (Illumina, San Diego, CA, USA). Updated coordinates of these SNPs in accordance with latest pig assembly can retrieve from http://www.animalgenome.org/repository/pig/. In various earlier mapping studies, this assay was employed to reveal the list of QTL regions, candidate genes and SNPs marker in association with different economically important traits in the pig by means of genome-wide association studies (GWAS) approach. For instance in boar taint, there were several independent QTL mapping studies done with the help of PorcineSNP60 on population of geographically distinct pigs (Zadinová, Kateřina, et al., 2016). A list of trait linked QTLs in pig can obtain from http://www.animalgenome.org/cgi-bin/QTLdb/SS/index. Another high density pig SNP assay is now accessible on market from GeneSeek, a Neogen company and contains almost 70,000 SNPs with mostly higher minor allele frequency markers distributed uniformly across entire pig genome. Same company also launches the GeneSeek Genomic Profiler for Porcine (GGP-Porcine) low density BeadChip which consists in nearly 8,500 SNPs for high density imputation (http://www.neogen.com/Agrigenomics/pdf/Slicks/GGP_PorcineFlyer.pdf). These two assays comprises genetic markers that could be directly associated with disease and performance traits including fat content and meat quality traits. The annotation of these SNPs is possible to obtain at http://www.animalgenome.org/repository/pig/Pig70K_SNP_annotations.csv.gz.

After identification of boar taint associated QTL regions and probable candidate genes, whole genome re-sequencing of multiple individuals can be followed to dissect QTLs regions in order to achieve fine mapping that lead to either narrowing of QTL region, candidate gene, functional mutation or haplotypes. This approach required multidisciplinary skills in bioinformatics, omics, and genetics. Data analysis part of sequencing experiment begins from the quality checking of either paired-end or single-end reads followed by the removal of duplicate reads, adaptor bases, low quality bases or very short reads. After filtering step, mapping of reads to reference genome and then downstream process will be followed. Downstream process is varied according to objective of the experiment. Typically in re-sequencing project, variants will be identified genome-wide, exome-wide or region-wide in multiple samples simultaneously. In case, where SNPs called for dissecting a specific trait, there is need to sort out high quality SNPs after estimation of their functional effect to perform high-throughput genotyping in certain number of individual samples and then successfully genotyped SNPs will be tested for associations with examined trait.

In one of the recent boar taint studies, whole genome sequencing of 55 animals including European and Asian pig breeds was performed to find out SNPs within regulatory regions of six differential candidate genes (*SPACA4, SYNGR4, TULP2, FTL, GLTSCR1* and *SULT2A1*) identified by using mRNA sequencing, and within coding region of all of the possible candidate genes located on porcine chromosome 6 (SSC6) in position between 48,585,961 bp and 52,336,598 bp (according to *Sscrofa10.2* assembly) underlying androstenone in fat tissue (Hidalgo, André M., et al., 2014). From this data, they identified single SNP (C/G at position 49,110,873 bp) reside inside a CpG island (49,110,687 bp – 49,110,889 bp) that could play important role in differential expression of *SULT2A1* gene (Hidalgo, André M., et al., 2014). This *SULT2A1* gene encodes the sulfotransferase enzyme to perform sulfoconjugation of α-androstenone in testis. This particular finding also points toward significance of CpG methylated regions in gene expression analysis. In the future, mapping of the methylated regions within QTL regions located on different pig chromosomes using bisulfite sequencing will be attractive practice to comprehend how they epigenetically regulate genes associated with boar taint.

A peak of genome-wide significant SNP effects on androstenone on SSC5 (P⌷>⌷6.8E-07) explaining 4% of the phenotypic variation in one recent study (Rowe, Suzanne J., et al., 2014). In same study, most significant GWAS result is found for *ASGA0025097* placed ~4 Mbp distal to another important *H3GA0016037* SNP on SSC 5. The genes of interest within this 4 Mbp region comprise the retinol dehydrogenase 5 (RDH5) and retinol dehydrogenase 16 (RDH16) genes. Another 17-beta hydroxysteroid dehyrdrogenase gene (HSD17B6) is located about 0.5 Mbp upstream of the *ASGA0025097* SNP. The 4 Mbp long regions between the two top SNPs is enriched with genes and showed high levels of LD in the Danish Landrace population studied. Oddly, many of the genes in this region encode olfactory receptors. The minor allele frequency for both SNPs (*ASGA0025097*, *H3GA0016037*) was 0.14 and the r^2^ between them was 0.68(Rowe, Suzanne J., et al., 2014).

Another study in which whole genome re-sequencing of 23 Duroc and 22 Landrace Norwegian male pigs was performed with an aim to ascertain a large numbers of SNPs and haplotypes underlying boar taint compounds within three QTL regions (identified based on large scale genome-wide association and LDLA mapping study; Grindflek E et al., 2011) on SSC 5, 13 and 7 underlying androstenone and/or skatole (Rahul et al., 2013, Rahul 2013).

So far in SSC5, an extensive and significant haplotype block of 69 SNPs, out of overall 132 SNPs, (significantly related with androstenone levels in fat (p-value <0.001)) for fat androstenone levels in Norwegian Duroc boars identified in which three plausible interesting candidate genes - RDH16/SRD9C8, short chain dehydrogenase/reductase family 9C, member 7 (SDR9C7) and HSD17B6. Phenotypic variance explained by each non-synonymous SNPs and 5’UTR SNP of candidate genes within this block is > 5 %. One of the haplotype blocks with frequency 0.66 is in association with considerably lower levels of androstenone. A single non-synonymous SNP in *RDH5* is found associated with androstenone fat (Rahul, Manuscript in Preparation) and in SSC 13, a total of 66 new and 15 Illumina genotyped SNPs used for assessing associations with the levels of androstenone in fat tissue. The association study shown that altogether 5 new SNPs and an known SNP of sodium channel, voltage-gated (SCN) genes significantly affecting the levels of fat androstenone (p<0.05) and with these 6 SNPs, a significant (p<0.05) haplotype block of 23-SNPs was constructed (Rahul, Manuscript in Preparation).

### RNA Sequencing

Quantitative and qualitative estimation of transcripts in response to different experimental conditions using RNA sequencing is now most widespread method for expression profiling in livestock and plant species. With RNA sequencing, it is also possible to discover variants, novel transcripts, and isoforms, splice sites, alternative transcription start sites, gene fusions and perform strand specific annotation of transcripts. Analysis of RNA sequencing data is based on presence and absence of reference assembly of an organism. RNA sequencing definitely provides clear advantage over genome sequencing where reference assembly not available. In absence of reference assembly, short reads obtained from mRNA sequencing can be *de novo* assembled into smallest set of transcripts to proceed to subsequent steps. Otherwise, analysis process is as same as DNA sequencing till read mapping except exclusion of duplicate reads to avoid the errors in estimating transcript abundances.

Recent two reference-based RNA sequencing studies on extreme levels of boar taint compounds have revealed a large number of candidate genes and SNPs in testis and liver of crossbreed German boars. In the case of androstenone, there were 46 genes found differentially expressed in testis while in the liver the numbers of differentially expressed genes were 25. Some important candidate genes revealed are *DKK2, DSP, CYP2B22, IP6K1, IRG6, HIST1H4K, MX1, IFIT2, CYP7A1, FMO5, KRT18, TSKU* and *HSD17B2*. The results of association mapping between SNPs and expressed genes have shown that SNPs in *IRG6, DSP,* and *IFIT2* genes were only related with low androstenone levels in testis while SNPs in *FMO5, HIST1H4K* and *TSKU* specifically linked with low androstenone levels in liver (Gunawan, Asep, et al., 2013a). While, in the case of skatole, they found 448 differentially expressed genes among those SNPs in *ATP5B, KRT8, PGM1, SLC22A7* and *IDH1* genes were found to be highly significant during association study of these SNPs in relation to skatole levels in boar liver (Gunawan, Asep, et al., 2013b). Moreover, exons of three genes (*ATP5B, KRT8,* and *PGM1)* were found to be differentially expressed during expression study of exon usage of these genes (Gunawan, Asep, et al., 2013b). An association study between polymorphisms of six genes and boar taint related compounds androstenone, skatole and indole in a boar population (n = 370) revealed that FMO5, CYP21 and ESR1 gene variants might have effects on the boar taint compounds (Neuhoff, Christiane, et al., 2015). Further, mRNA expression study revealed that levels of CYP21 and ESR1 were found higher in castrated boar, in relation to non-castrated boars; while, the expression of FMO5 and ESR1 were found higher in low boar taint, in comparison to high boar taint in liver tissue(Neuhoff, Christiane, et al., 2015).

Male pigs (n=48) with low, medium and high genetic merit of boar taint were selected for transcriptomic profiling by RNA-Seq revealed more than 80 deregulated genes that functionally categorized to Gonadotropin releasing hormone receptor and Wnt signaling pathways, linked with reproductive maturation and proliferation of spermatogonia, respectively (Drag, Markus, et al., 2016). Real-time polymerase chain reaction experiments on 26 testes and 12 adipose tissue samples from pubertal boars using 21 genes unraveled that only transcriptional levels of CYP17, CYP11A1, CYP19A, SRD5A1, and AKR1C-pig6 are correlated with the fat concentration of androstenone (0.57 < r < 0.70, P <0.05) suggesting that the levels of androstenone in fat is interrelated to the production in testes of androstenone and more generally to all sex steroids (Robic, A., et al., 2016).

The role of two micro RNA- *miR-122* and *miR-378* investigated in liver of Jinhua pig and reported that these two miRNAs have associations with three previously reported candidate genes-S*ULT1A1, SULT2A1* and *CYP2E1*.(YiTao, Ma, et al., 2013). Hence, it will be essential to consider the role of small RNAs in the regulation of gene activity in boar taint. Specifically, miRNA is known to degrade transcripts of genes to control their expression, therefore, expression profiling of miRNAs using protocol of small RNA sequencing in boar testis and liver can provide the list of miRNAs that can epigenetically control the activity of several boar taint associated genes together. A list of annotated pigs small RNAs can retrieve from URL: http://rth.dk/resources/rnannotator/susscr102/ and further role of and the latest pig specific array based on miRBase version 21 and features all 407 unique mature miRNAs available at (http://www.lcsciences.com/discovery/applications/transcriptomics/mirna-profiling/mirna/mirna-availablearrays/pig-array-content/).

Estimation of copy number variations (CNVs) from either exome/targeted/whole genome sequencing would be helpful in understanding their direct effect on gene expression, and on machinery of smallRNAs (Avinash, M. V., et al., 2014, Persengiev, Stephan et al., 2014). In Pig, there is one study in which detection of CNV regions (CNVRs) was performed by using Porcine SNP60 BeadChip data on autosomal chromosomes using a pedigree from an Iberian x Landrace (IBMAP) cross (Ramayo-Caldas, Yuliaxis, et al., 2010). Overall 49 CNVRs were found in 13 autosomal chromosomes and these CNVRs exhibited Mendelian inheritance across 427 individuals belonging to several generations of an Iberian x Landrace cross. CNVs were reported in many pig breeds as mentioned in (Xie, Jian, et al., 2016). Thus, CNVs can be narrow down to boar taint -genes, -miRNAs and/or -variants in order to comprehend their role and then use them as genetic marker for boar taint selection.

### Interactomics and Metabolomics

Analysis of high-throughput sequencing data gives a list of plausible candidate genes, differential transcripts and biomarkers underlying a given trait, but whole biological mechanism thats drive the interplay between genes, protein, RNAs or biomarkers would not be understood without deciphering the pathways and interactions that link them. The process of decoding and analyzing comprehensive set of molecular networks in a given organism, tissue or cell is termed as *interactomics*. From RNA sequencing data, it is possible to import the list of differentially expressed genes into pathway and interaction prediction tools including Ingenuity Pathway Analysis (IPA) software, Cytoscape, DAVID, Omiras and GSEA that provide all of the putative networks and the metabolic pathways affecting a given trait. Latest interactomics study in boar taint reported a total of 1023 significant interactions between 826 genes weighted with Pearson correlation coefficients for extreme levels of androstenone in German boars and altogether 92 pathways enriched for these interactions using gene expression data from previous RNA sequencing experiment done by the same author (Sahadevan, S, et al, 2015). This study also stated that glutathione metabolism, sphingolipid metabolism, fatty acid metabolism and cAMP-PKA/PKC signaling were foremost pathways that regulate steroid and androstenone biosynthesis in higher and lower androstenone level boars (Sahadevan, S, et al, 2015).

Metabolomics study of samples from a particular tissue can identify a set of metabolite biomarkers that distinguish two different conditions for instance, metabolomics data can be used to reveal additional biomarkers that differentiate the adipose tissue from non-tainted and tainted boars. Non-targeted metabolomics study is pursued to recognize the entire potential previously unknown biomarkers or metabolites ion within certain mass range. Previous liquid chromatography mass spectrometry (LC-MS) based non-targeted metabolomics study in boar taint had identified 15 unidentified compounds and three identified compounds- testosterone, androstenadione and 3-oxohexadeanoic acid (Olson M et al., 2012). These 18 compounds were able to explain the significant difference between untainted and tainted samples taken from adipose tissue of all 32 pigs. All the data analysis was done in Progenesis CoMet software (http://www.nonlinear.com/progenesis/qi/). Skatole levels in fat is found negatively correlated to CYP2E1 activity and positively to 3-hydroxy-3-methyloxindole (HMOI), indole-3-carboxylic acid (ICA) and 2-aminoacetophenone in urine using liquid chromatography-MS of samples from urine (n=46), blood (n=12), liver (n=25) and adipose tissue (n=46) samples prepared from a total of 46 entire male pigs (Brunius C et al., 2016). In forthcoming days, studies combining GWAS and metabolomics profiling data (mGWAS) can be worthwhile to pursue in order to find out individualistic genetic regulation of metabolites in non-tainted and tainted boars.

### Integration of omics data

Utilization of different kinds of high-throughput omics data in dissecting boar taint trait in intact male pigs will not be rewarding until there is an effort to integrate outcomes coming from different omics studies. Defining entire biological mechanism of a trait with one state-of-the-art functional genomics approach is a daunting task. It is critical to decipher complex interactions exist between different molecules in adipose, testicular and liver tissues to understand overall mechanism behind the occurrence of boar taint. However, this approach needs professionals with broad range of skills in bioinformatics, genetics, mathematics, statistics, molecular biology, and chemistry and computer science. For instance, recently launched GenSAP project (http://gensap.au.dk/), which aims to implement and automate genomic selection program in different livestock and crop species with consistent improvement in its implementation through input coming from integrated data of whole genome sequencing, functional genomics, epigenomics, and complex phenotyping technologies. And also there is alternate way to encourage integration of data is to share existing pig related omics data by storing them in open source database or cloud storage platforms so even individuals from non-biological disciplines can access these data and make best use of their data mining knowledge to establish, and interpret the links between various omics data.

## Conclusion

There is no doubt that deluge of omics data and bioinformatics resources has completely transform the manner with which animal breeding and selection practices pursued earlier. Definitely in future, more pig based omics data will likely be generated to enrich an existing resources with excess of genetic information to continuously improve the method of genetically selecting the entire male pigs against increased boar taint levels. Integration of distinct omics data would be possibly plays a key role in solving boar taint problem to a greater extent.

